# BlotIt - Optimal alignment of western blot and qPCR experiments

**DOI:** 10.1101/2022.02.09.479689

**Authors:** Svenja Kemmer, Severin Bang, Marcus Rosenblatt, Jens Timmer, Daniel Kaschek

## Abstract

Biological systems are frequently analyzed by means of mechanistic mathematical models. In order to infer model parameters and provide a useful model that can be employed for systems understanding and hypothesis testing, the model is often calibrated on quantitative, time-resolved data. To do so, it is typically important to compare experimental measurements over broad time ranges and various experimental conditions, e.g. perturbations of the biological system. However, most of the established experimental techniques such as Western blot, or quantitative real-time polymerase chain reaction only provide measurements on a relative scale, since different sample volumes, experimental adjustments or varying development times of a gel lead to systematic shifts in the data. In turn, the number of measurements corresponding to the same scale enabling comparability is limited. Here, we present a new flexible method to align measurement data that obeys different scaling factors. We propose an alignment model to estimate these scaling factors and provide the possibility to adapt this model depending on the measurement technique of interest. In addition, an error model can be specified to adequately weight the different data points and obtain scaling-model based confidence intervals of the finally scaled data points. Our approach is applicable to all sorts of relative measurements and does not need a particular experimental condition that has been measured over all available scales. An implementation of the method is provided with the R package blotIt including refined ways of visualization.

## Introduction

The approach of mathematical modeling to analyse and understand dynamic processes of biological systems requires the collection and quantification of time-resolved experimental data for many different experimental conditions [1–4]. Frequently, the generation of these type of data is achieved by techniques like Western blotting [5,6], quantitative real-time polymerase chain reaction [7], reverse phase protein arrays [8] or flow and mass cytometry [9] which only generate measurements on a relative scale. Therefor, the number of experiments that are comparable to each other, i.e. provided on the same measurement scale, is typically limited by the experimental setup, which constitutes a bottleneck for mathematical models with high complexity.

In the following we focus on Western blotting as a well-established and commonly used technique. In this technique, protein abundances are measured on a relative scale by chemiluminescent antibodies binding to the respective proteins embedded in a gel. Let us consider such a time-course experiment that has been performed twice and quantified with Western blot. The experimental setting is assumed to be the same between the two experiments, i.e. the same biological or experimental conditions were measured, but on two different gels. Since the Western blot technique only provides a relative measurement, the obtained data points presumably show a similar dynamical behavior. However, they do not coincide with each other in absolute numbers due to experimental errors and different measurement scales. The corresponding unknown scaling factor, i.e. the ratio between the measurement scales, can be inferred relatively simply by aligning both measurement profiles to each other.

Now, let us consider another experiment where time courses of two different experimental conditions, as for example stimulation doses, have been measured separately on two different Western blots. Here, it cannot be distinguished whether the difference in the results is occurring due to the experimental condition or due to the different measurement scales. In particular, the scaling factor between the two blots cannot be estimated in this case.

One way to circumvent the missing comparability between measurements is to add recombinant proteins to the Western blot samples allowing for an absolute-scale quantification [10]. However, this approach is very expensive and time-consuming and scales with the number of measured proteins [11]. In Systems Biology, where the data is employed to estimate parameters of a mathematical model, it is a common approach to determine the scaling parameters of the different blots together with the remaining parameters of the model [12]. Besides the disadvantage of enlarging the parameter space drastically, the estimates of the scaling parameters might be biased by the model equations hampering hypothesis testing and therefor interpretation of the results [13].

As a generally applicable alternative, the Western blot experiments can be designed in a way that a certain experimental overlap exists between different blots, meaning that the same experimental condition is measured multiple times. Degasperi et al. [14] present a method to analytically determine the corresponding scaling factors based on such data. However, to be able to apply this method, there needs to be at least one experimental condition that has been measured on all blots which implies additional planing effort, might be limited by the availability of the overlap sample and complicates the use of experiments performed at a later time point.

Here, we present a new data-based approach for the estimation of scaling parameters which is also applicable in the absence of a unique condition overlapping across all available scales. It is sufficient when the independent experiments are connected by pairwise overlapping conditions. In addition, the implemented method provides not only the possibility to obtain scaling parameters and therefor align data points of different Western blots but also to compute confidence intervals for the results by applying a user-defined error model [15]. We implemented this method in the R package blotIt.

## Methods

When analyzing the measured values of a hypothetical experiment, we define three classes of effects: (I) *Biological effects* describe biological conditions as for example different targets, stimulation doses, inhibition treatments or measurement time points of a dynamical process. For each unique set of biological effects *i*, there exists one *true value y_i_*. (II) *Scaling effects* describe the systematic influence of the measurement techniques and evaluation routines on the particular numerical value that is obtained. In the example of Western blotting, these scaling effects include for example development time, sample loading or gel thickness. All scaling effects add up to a scaling factor *s_j_* that equally affects the measurements of all *y_i_* within the respective experiment. Only measurements that underlie the same scaling factor can *a priori* be considered as comparable. In other words, repeated experiments measuring the same effect *y_i_* result in a set of values *Y_ij_*, where the indices imply that *y_i_* is affected by experiment-specific scaling factors *s_j_*. Some properties, e.g. gel imperfections, do not affect the whole experiment uniformly, but neighbouring lanes can be influenced by a systematic error. To resolve this, randomized sample loading is advised [16], ensuring that the resulting errors are independent. (III) *Residual noise*: In addition to the systematic error sources (I-II), each measurement *Y_ij_* is affected by stochastic noise _*ij*_. Based on these three error sources, we present in the following an approach to align the numerical values of different experiments with the aim of retrieving one comparable data set.

### Definition of the alignment model

In mathematical terms, the influence of scaling factors *s* on true values *y* is described by

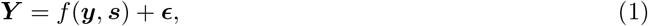

where *f* (***y***, ***s***) is the scaling model and reflects the noise of the measurement which is assumed to be normally distributed with 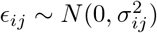, where *σ_ij_* is the standard deviation of the normal distribution, and the indices imply that each measurement *Y_ij_* can in principle have its own error distribution. To assess the individual error distribution, an error model *h* is introduced

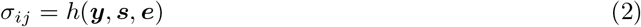

which is based on the error model parameters ***e***. This approach of estimating the errors is superior to the calculation of errors based on the spread of measurement values, as the error model considers the variance information from all experimental data and thus allows for a reliable error estimate even for conditions with small numbers of replicates.

When considering Western blot data as a typical use case for the here presented method, the model simplifies to a scaling model with gel-dependent scaling effects. The equation then reads

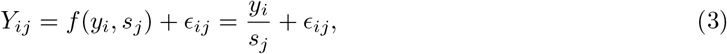

where the measurement *Y_ij_* corresponds to the true value *y_i_* affected by the scaling factor *s_j_* and the noise *ϵ_ij_*. It is known that the error of Western blot data can be described by a relative error model [15]. In this case the standard deviation for the measurement *Y_ij_* is estimated by

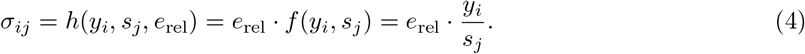

In this particular error model, there is only one error parameter *e*_rel_ for all measurements, from which the measurement errors *σ_ij_* are inferred by multiplying *e*_rel_ with the corresponding model evaluation. Note that more complex error models including also absolute contributions or gel dependent effects could be chosen.

Depending on the measurement technique, the data could be given on the logarithmic scale, and the error model *h* has to be adjusted accordingly. This is the case e.g. for qPCR data, which is typically provided on log_2_ scale:

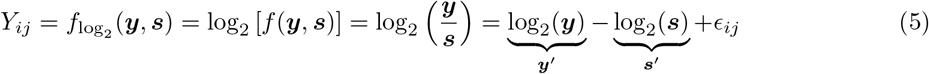

Here, the relative error model becomes an absolute one:

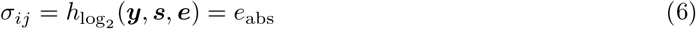

Together, Eq (1) and Eq (2) describe a combined scaling and error model formulation based on the assumption that true values of measurements are influenced by scaling factors and experimental noise. All parameters ***y***, ***s*** and ***e*** are *a priori* unknown and have to be determined based on the data.

### From the original to a common data scale

Evaluating the data with the presented method results in three representations of the data, each of them with its own meaning and application in different contexts: (i) The *scaled* data representation contains the original replicate data transferred to the common scale, (ii) the *aligned* data representation reflects the underlying estimated true values, and (iii) the *predicted* value representation corresponds to model evaluations back on the original scale. The alignment process as described in the following is schematically visualized in Fig 1.

**Fig 1.**
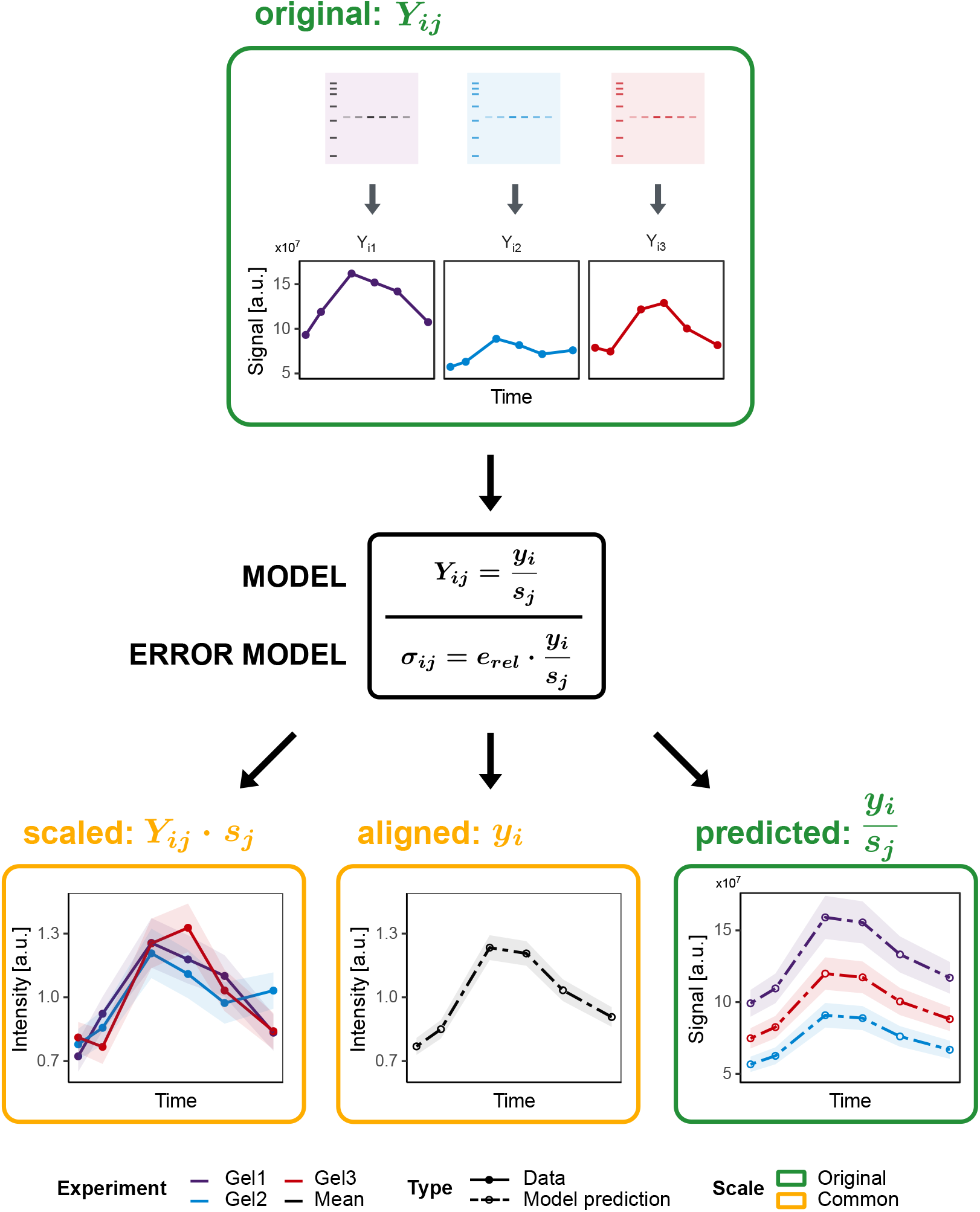
Overview of the blotIt alignment procedure. Top: Exemplary raw data on the original scale (original) as being measured on three different blots indicated by the color. Middle: Raw data is fitted by the alignment model to estimate scaling parameters *s_j_* and the underlying true values *y_i_*. Error parameters *e_rel_* are simultaneously estimated by means of an error model. Bottom: The procedure outputs three different ways to visualize the result: (I) Single replicates aligned to the common scale (scaled), (II) the time course of estimated true values (aligned), and (III) a prediction for the replicates on the original scale (predicted). Uncertainties are shown as shaded areas.

Initially, the numerical values of each experiment are on their own *original scale* shown by three representative example time courses at the top of Fig 1. In particular, these measurements are not comparable to each other. Now, let us for the moment assume that an optimal set of parameters 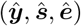 has been found. With these parameters, we define a *common scale* as the scale on which all measurements shall be directly comparable. The model assumes that all true values ***y*** are on this common scale, and describes how the scaling has to be applied to a true value *y_i_* to match the respective measurement *Y_ij_*. To retrieve the true values *y_i_* form a respective measurement *Y_ij_* under the scaling *s_j_*, the *inverse* of this model has to be evaluated

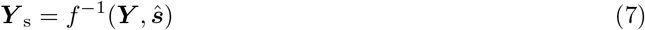

with estimated scaling parameters 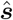 and measured data ***Y*** on the original scale. The resulting data set with replicates aligned to the common scale is thus called *scaled*, as shown on the lower left in Fig 1. The scaled data set still contains the information about each independent experiment, but all measurement values are directly comparable. This is useful for the comparison between experiments, e.g. to determine potential outliers and identifying experiments with obviously significant measurement errors. As input for a dynamical modeling approach, the scaled data set is preferred in comparison to the original data as it is already on a common scale. Thus, individual scaling parameters do not need to be determined by the dynamical model, thereby decreasing the parameter space. In some occasions, the mean values along with their estimated errors are favored over the scaled replicates. This might be the case, e.g., when only small numbers of replicates are given. In this case, the *aligned* data set can be used, containing the estimated true values 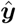 along with the corresponding estimated errors 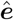. These estimated errors might be a more reliable description of the data spread compared to the information provided by a small number of replicates.

To identify discrepancies between the true values determined based on all measurements, and the measurements of a single experiment, the true values can be scaled back to the original scale. This new data set is termed *predicted* and consists of the direct alignment model evaluations using the estimated true values and scaling parameters:

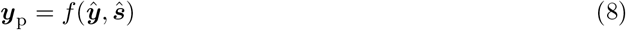

Note that this data set is again on the original scale, and thus does not provide comparability between the experiments.

### Parameter estimation

The presented method brings measurements from different experiments to a common scale applying an alignment and an error model. Assuming the parameters obey Gaussian statistics, the best maximum-likelihood estimate 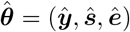 is the set of parameters, for which the spread of replicates is minimal [17]:

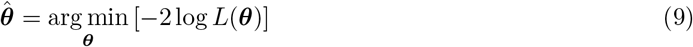

Here, the log-likelihood function log *L*(***θ***) consists of three terms:

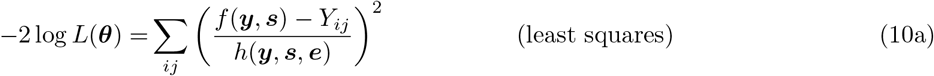

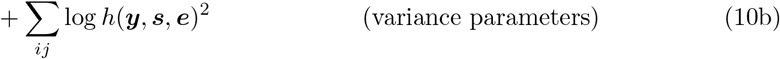

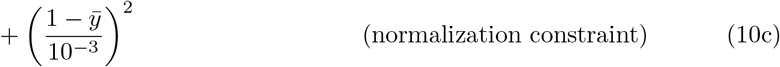

The special form of the log-likelihood function presented in Eq (10a) and Eq (10b) is based on the assumption that observations are affected by normally distributed residual noise. The first term (10a) includes the weighted least squares, namely the difference between model prediction *f* (***y***, ***s***) and corresponding measurements *Y_ij_*. Residuals are weighted by the variance 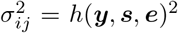. The second term (10b) accounts for the simultaneous optimization of the error model *h*(***y***, **s**, ***e***) and thereby estimation of error parameters. While the first two terms ensure minimization of the spread between experiments, the third term (10c) forces the mean of the estimated true values 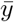 to be one during the optimization process and thereby introduces the common scale.

For computational reasons, the parameters are per default transferred to logarithmic scale prior to the estimation. This drastically improves numerical stability especially when the input data varies over multiple orders of magnitude. Estimated parameter values are subsequently transformed back and reported on the linear scale.

### Error determination

In Fig 1, we introduced different output data representations as result of the alignment model. As a major advantage of this formulation, a measure of uncertainty, i.e. a statistical error, can be determined for each of these data sets, as summarized in Table 1 and explained in the following.

**Table 1.**
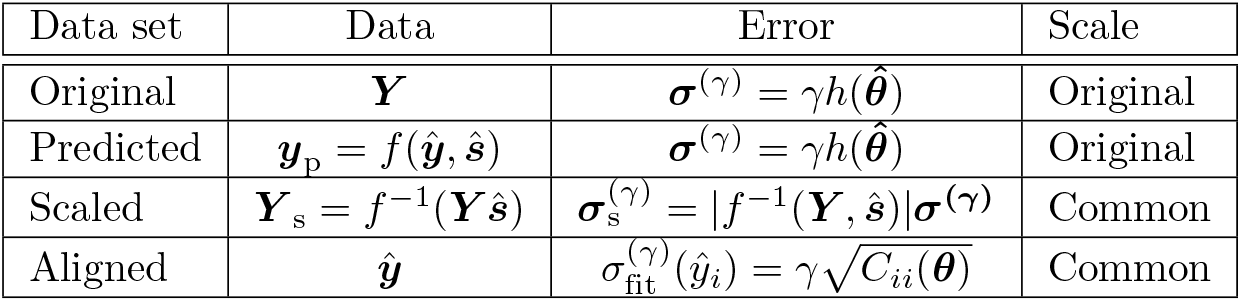
Overview table of the different output data sets of blotIt.

First of all, the number of estimated parameters *n*_P_ = |***θ***| in the alignment model is typically quite high compared to the number of data points 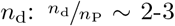. Under such conditions, the maximum-likelihood estimation tends to underestimate the standard deviation in a sample. The effect is more apparent for small samples or, equivalently, when many parameters are estimated from few data points. We account for this bias by applying Bessel’s correction to scale the estimated sample standard deviation by a factor

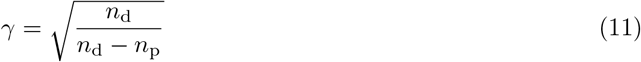

that is multiplied with the estimated standard deviation of the measured data within blotIt:

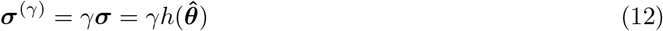

The error model is evaluated on the scale of the original observations. To retrieve the error of the scaled data, Gaussian error propagation is employed:

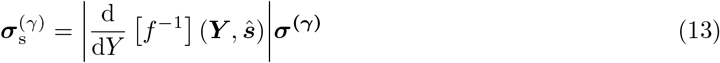

While all errors considered until now quantify the uncertainty of the measurement, the error of the estimated true values 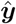 are calculated qualitatively differently. Since 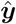 are model parameters, their errors have to be estimated by the model uncertainty itself. In maximum likelihood estimation, the uncertainty is reflected in the local curvature of the likelihood landscape around the determined parameter value. The uncertainty of the *l*-th fitted parameter 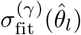 is given by

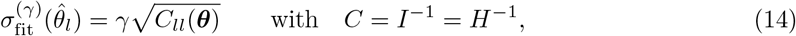

where the local curvature is approximated by the square root of the *ll*-diagonal element of the covariance matrix *C* given by the Fisher information matrix *I*, which itself is represented by the Hessian *H* that is calculated during the optimization process. Note that the Bessel correction *γ* is applied here, too.

To sum up, the blotIt approach provides four measures of uncertainty (Table 1). Uncertainty provided with the original data set is estimated based on the error model. This error is reflective of the between-replicate variability and, as such, is comparable to the standard deviation of a single measurement. Uncertainty provided with the predicted data set is the same as for the original data set and, thus, reflective of the standard deviation of the single measurement. The uncertainty provided with the scaled data set is the error of the original data set translated to a different scale, i.e. the common scale. Also this error is reflective of the standard deviation of the single measurement. Finally, uncertainty provided with the aligned data set is the estimation uncertainty of the estimated *true* concentrations meaning that this error is comparable to the standard error of the mean. Therefor, with more and more replicate measurements, the error of the aligned data set becomes smaller, whereas the errors provided with original, predicted, and scaled data sets consolidate.

## 1 Applications

In the following, the application of blotIt is illustrated by means of a published data set [4] that provides time-resolved measurements of the phosphorylation of cytoplasmic *Signal Transducer and Activator of Transcription 1* (STAT1) and mRNA levels of the *Suppressor of Cytokine Signaling1* (SOCS1). Both targets are involved in the Interferon alpha (IFNα) signaling pathway, where STAT1 acts as a transcription factor regulating, among others, SOCS1 expression. Three different IFNα concentrations were used to induce signal transduction and thereby phosphorylation of STAT1 as well as expression of SOCS1. Phosphorylation dynamics of cytoplasmic STAT1 were quantified by Western blot experiments, while SOCS1-mRNA levels were measured by qPCR. The three IFNα doses used for stimulation correspond to three experimental conditions that are distinguished in the following. The scaling and thereby alignment of the Western blot as well as the qPCR data, derived from three experiments each, are performed with the R package blotIt.

### Alignment of Western blot data

Cytoplasmic STAT1 protein was quantified in three different experiments. Measurements are thus only available on different scales as they originate from distinct gels. Therefore, one can not directly judge the dynamics of the final time course from investigating the raw data (Fig 2a). Moreover, a direct comparison for example between experiment two and three is not possible, since these have not been measured together on the same gel. Instead, as shown in Fig 2b, an overlap exists between experiments 1 & 2, and 1 & 3 by two replicates, respectively. This allows the alignment of the three experiments and enables a comparison between all replicates. If no overlap existed between gels, it would not be possible to determine or define a common scale. In this case, blotIt would determine scaling factors separately, and the resulting scaled values would not be comparable between experiments.

**Fig 2.**
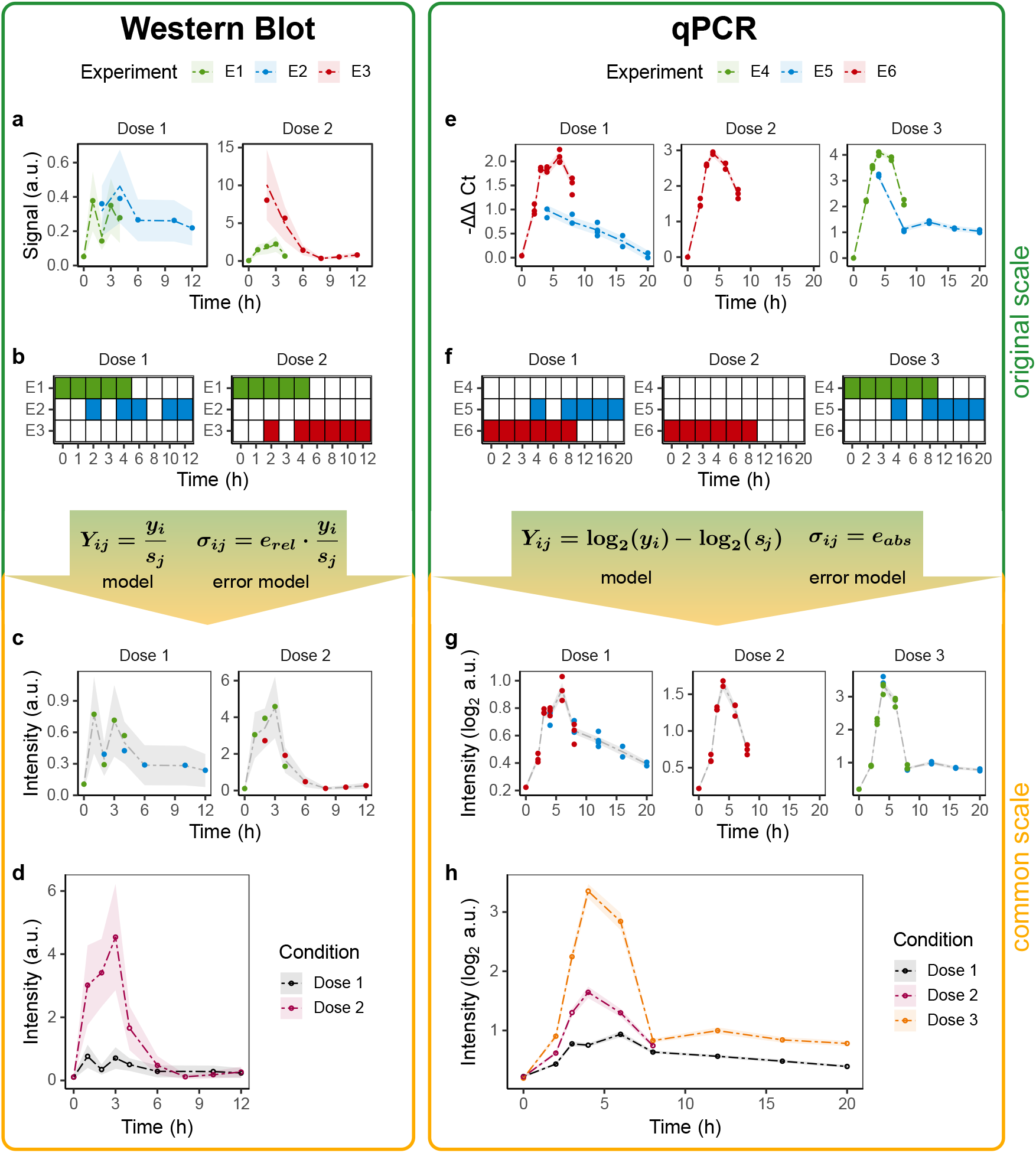
Application example for the alignment of Western blot and qPCR data. Raw data of cytoplasmic pSTAT1, measured by Western blot, and SOCS1 mRNA quantified with qPCR, was taken as a subset from [4]. (a, e) Raw data is shown on the original scale (dots) compared to the predictions (dashed interpolating lines) as output by the model. Color indicates the different experiments (gels). (b, f) Illustration of the experimental overlap. Rows correspond to the different gels (*scaling effects*), columns correspond to different experimental conditions (*biological effects*). Tiles indicate whether the respective conditions was measured (colored) on the respective gel or not (white). (c, g) Data points after the alignment are shown. Scaled replicates (dots) are colored according to their original gel. On the same common scale, estimated true values are shown as gray interpolating lines. (d, h) Aligned data (dots) and trajectories (linearly interpolating lines) are depicted on the common scale. Color indicates the experimental condition.

Within blotIt, scaling of these Western blot data is performed via the alignment function `alignReplicates()` called as

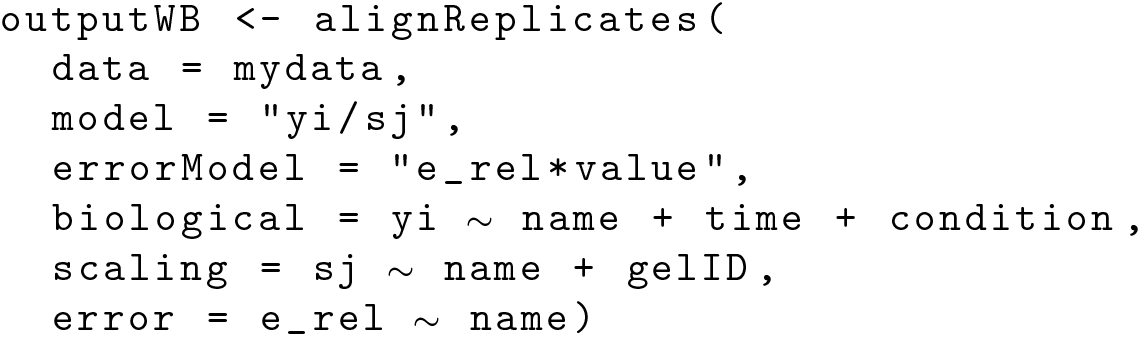

allowing an individualized structure of model and error model. The structure of the input data set is oriented towards a recently developed data sharing standard for dynamic modeling, in particular the measurement file of PEtab [18]. It has to be provided as a `data.frame` with obligatory columns `name, time` and `value`, specifying the observed target, the measurement time and the measurement value. Further columns characterize additional biological effects, which have to be distinguished, and scaling effects, e.g. the experimental `condition` or the `gelID` of the Western blot.

The alignment function allows to define these effects as a function of *y_i_* or *s_j_* in an additive manner. `name` and `time` are obligatory for the argument biological, as different targets at different time points are independent and have to be distinguished. The scaling argument requires the parameter `name`, as every observed target may scale differently. Also the error, here `e_rel`, can be specified. If the same value of `e_rel` should be assumed for all conditions, the error is only specified by the target, i.e. `name`.

Alignment of the pSTAT1 data brings the measured data points from different experiments to a common scale. The `output` of `alignReplicates()` is a list with the entries *scaled*, displaying the scaled replicates *Y_ij_ s_j_* (Fig 2c) and *aligned*, containing the estimated true values *y_i_* (Fig 2d), both with uncertainties. Further listed elements describe the original data, data predictions based on the alignment model, and the respective estimated scaling parameters.

### Alignment of qPCR data

The flexibility of the model based alignment approach allows to process qualitatively different data with only minor adjustments. Thus, in addition to the linear Western blot data, also quantitative real-time polymerase chain reaction (qPCR) data can be analyzed with blotIt. During the process of mRNA quantification in qPCR measurements, a small region of the mRNA of interest is amplified in a sequence of replication cycles. The mRNA concentration is therefor measured in Cycles to Threshold (*C_t_*) of PCR, a relative value that represents the cycle number at which the amount of amplified DNA reaches a defined threshold level. This threshold is in general individually chosen for each experiment. Since the amount of mRNA is approximately duplicated in each cycle of the PCR, the *C_t_* value is on the log2 scale. The inferred quantity Δ*C_t_* describes the difference in cycles between the target and a reference gene, where the reference gene can be a *housekeeper* which is known to remain relatively stable in response to any treatment. To assess the dynamic development of mRNA expression the ΔΔ*C_t_* can be used, representing the difference in Δ*C_t_* between the target and a reference condition [19]. Here, an intuitive reference condition is the zero time point. Because higher *C_t_* values correspond to lower mRNA abundance in the sample, the quantity ΔΔ*C_t_* is used for the description of expression dynamics.

The freedom of choice for the detection threshold results in an experiment-specific shift in the number of cycles until detection. Together with the offsets introduced in the ΔΔ*C_t_* calculation by potential measurement variances of the zero time point and the housekeeper genes, this can be summarized into one offset parameter. Since the data is of logarithmic nature, this offset reflects a multiplicative scaling on the linear scale. The alignment model and the corresponding error model thus have to be log-transformed, which results in an additive model with an absolute error description. The `alignReplicates()` function call is adjusted accordingly:

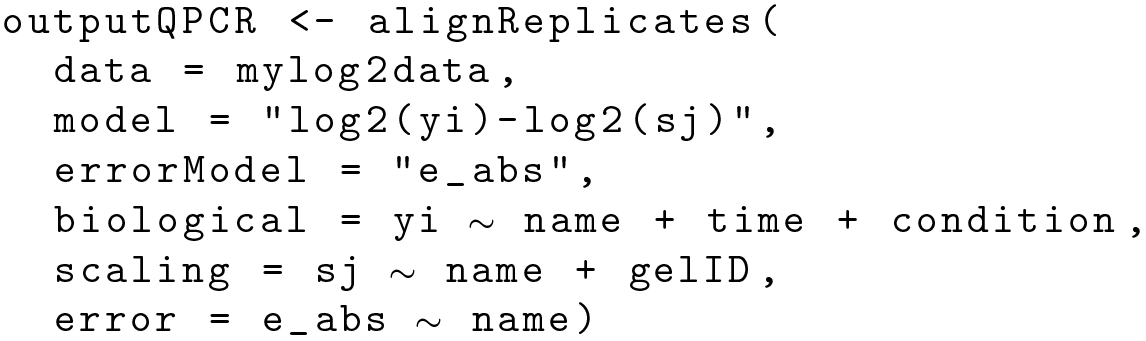

The results of the alignment process `outputQPCR` are analogous to those described for the Western blot data. They are described for the example data set of SOCS1-mRNA on the right hand side of Fig 2 (e-h). Data was quantified in three experiments, where experiment four and six are not comparable on the original scale and differences in conditions only become apparent on the common scale. Furthermore, the dynamics of the full time course are not visible at the level of the single experiments (Fig 2e) but can be analyzed after scaling in Fig 2h. Since the original scale is logarithmic this is also true for the common scale.

## Discussion

In many cases, biological data is generated in a way that does not allow a direct comparison between different measurements. Reasons can be differences in sample loading, antibody binding or discrepancies between various gels in the case of Western blotting. In turn, these artifacts lead to different measurement scales for the experimental data and mask the effects of biologically different conditions like treatments and measurement time points in dynamical processes.

Analyzing longitudinal data or dose response data without proper preprocessing is not possible when the measurements are affected by different scaling factors. We here present a method to scale data of independent experiments to one common scale, where the data is directly comparable. In addition to the *original* and the *scaled* data, we provide two further outputs of the algorithm: (1) *aligned* data, i.e. the *true values* obtained when the impact of different scaling and residual noise is removed; and (2) *predicted* data, i.e. the values on the original scale of the experiments obtained when only the impact of residual noise is removed.

Previously established strategies to correct for the scaling differences include the usage of recombinant proteins to transform each measurement to an absolute scale, or theoretical approaches to determine a common scale as discussed in [14]. In the latter, scaling factors are determined by analytically minimizing differences in scaling between the experiments. The here presented method follows a similar idea but determines the scaling factors via a more flexible numerical optimization. Our approach has the benefit that no single condition needs to be present within all experiments, but instead it is sufficient to have a pairwise overlap of measured conditions between different experiments. We are not only able to determine the scaling factors, but also estimate underlying true values, i.e. maximum-likelihood estimates for the true values disregarding the experimental scaling artifacts. Asymptotic confidence intervals based on the Fisher Information Matrix are provided for the scaling factors as well as for the estimated true values.

One typical field of application for biological time course data is dynamical modeling where it is often beneficial to reduce the number of estimated parameters to a minimum, e.g. via data preprocessing. By determining a common scale with the presented method, it is not necessary to include experiment-specific scaling factors in the model, decreasing the parameter space and therefore the model complexity. While the scaled data set consists of all replicates, the aligned data set represents the averages and the information about the spread of replicates is encoded in the corresponding errors.

By utilizing numerical optimization, alignment model and error model can be flexibly adapted to the appropriate scaling mechanism for the data at hand. One can therefore account for example for data on the logarithmic scale or apply customized scaling approaches. The same freedom applies to the error model with the benefit to individually include relative and absolute errors. With this flexibility, blotIt can not only be applied to data generated by Western blotting, but to all use cases where relative data is generated like quantitative real-time PCR, reverse phase protein arrays as well as flow and mass cytometry.

## Code Availability

The R package blotIt is available on GitHub at https://github.com/JetiLab/blotIt, where more information about system requirements and installation is available.

## Acknowledgement

We thank Marcel Schilling and Elisa Holstein for instructive feedback on the experimental procedure of qPCR measurements.

## Author Contributions

Conceptualized and implemented the methodology: DK. Developed the software: DK. Extended the software: SB, SK. Software maintenance: SB. Visualization: SK. Wrote the paper: SK, SB, MR. Reviewed the manuscript: DK, JT. Funding acquisition: JT.

